# Prediction of Protein Half-lives from Amino Acid Sequences by Protein Language Models

**DOI:** 10.1101/2024.09.10.612367

**Authors:** Tatsuya Sagawa, Eisuke Kanao, Kosuke Ogata, Koshi Imami, Yasushi Ishihama

**Author notes:** Correspondence and requests for materials should be addressed to Y.I.

## Abstract

We developed a protein half-life prediction model, PLTNUM, based on a protein language model using an extensive dataset of protein sequences and protein half-lives from the NIH3T3 mouse embryo fibroblast cell line as a training set. PLTNUM achieved an accuracy of 71% on validation data and showed robust performance with an ROC of 0.73 when applied to a human cell line dataset. By incorporating Shapley Additive Explanations (SHAP) into PLTNUM, we identified key factors contributing to shorter protein half-lives, such as cysteine-containing domains and intrinsically disordered regions. Using SHAP values, PLTNUM can also predict potential degron sequences that shorten protein half-lives. This model provides a platform for elucidating the sequence dependency of protein half-lives, while the uncertainty in predictions underscores the importance of biological context in influencing protein half-lives.

## Introduction

Protein abundances reflect a dynamic balance of various interconnected processes, including mRNA transcription, processing, and degradation, as well as protein translation, localization, modification, and programmed destruction. These processes are essential for maintaining the quality and quantity of protein pools within cellular homeostasis, and their dysregulation leads to various diseases^1–7^. Traditionally, mRNA abundance levels have been viewed as the primary determinant of protein abundance. However, the genome-wide correlation between mRNA and protein abundance levels is weak, with Pearson correlation coefficients ranging from 0.4 to 0.8^8,9^, underscoring the importance of post-mRNA regulatory mechanisms. Indeed, although mRNA abundance levels and ribosome density determine the protein translation rate, individual genes with similar mRNA levels can yield vastly different protein abundances^10^. Therefore, a comprehensive consideration of the impact of protein degradation is required for an understanding of protein abundance.

Recent advances in mass spectrometry (MS)-based proteomics have provided unprecedented opportunities for large-scale investigations of protein half-lives. Initially, protein half-life studies relied on pulse-chase radiolabeling^11–14^, a method that required stringent safety precautions and often resulted in disruption of homeostasis due to chemical treatment^15^. Alternative approaches, such as labeling proteins with fluorescent proteins or epitopes^10,16,17^, also have intrinsic limitations, including steric hindrance, photobleaching, and specificity issues, potentially failing to reflect protein half-lives accurately^15^. On the other hand, the combination of proteome analysis by LC/MS/MS with stable isotope labeling by amino acids in cell culture (SILAC) has overcome these disadvantages. SILAC enables a comprehensive global proteome assay of protein half-lives^18,19^ by measuring the time-dependent incorporation of “heavy” isotope-labeled amino acids into newly synthesized proteins to quantify synthesis rates. Simultaneously, a decrease in the sum of ‘light’ and ‘heavy’ amino acids is assessed at each pulse time point as an indicator of protein degradation. This dynamic SILAC or pulsed SILAC setup facilitates parallel measurement of the sequences and half-lives of thousands of endogenous proteins in cells, taking into account both protein synthesis and degradation. In parallel, over the past 30 years, significant advancements have been made in understanding protein degradation mechanisms, including the ubiquitin-proteasome system, autophagy-lysosome system and lysosome system. These advancements have led to the discovery of several principles governing the relationship between protein degradation and protein sequence. For instance, in the ubiquitin-proteasome system, the binding specificity of E3 ligases is partially controlled by short sequence motifs on substrates, known as “degrons,” typically less than 10 amino acids long^20–22^. The first identified degron was found at the amino terminus of a protein, leading to the formulation of the N-end rule^20^. Recently, the variety of degrons has expanded to include C-terminal degrons and degrons derived from internal sequences^23–26^. Furthermore, targeted protein degradation using degrons has emerged as a promising therapeutic strategy for regulating protein expression levels, potentially addressing “undruggable” components of the proteome. In this context, auxin-inducible degrons (AIDs) and proteolysis-targeting chimeras (PROTACs) have already demonstrated their utility in clinical settings^27–32^. Thus, understanding the relationship between protein degradation and protein sequence can also provide new insights into molecular design for drug discovery.

Natural language processing (NLP), a field that integrates linguistics and artificial intelligence, has made significant strides with the advent of deep learning and machine learning, leading to remarkable progress in areas such as sentiment analysis^33,34^, text summarization^35^, translation^36^, and text generation^37^. Protein sequences, analogous to human language, can be naturally represented as strings of letters, making NLP algorithms applicable to protein research^38–41^. In this context, protein language models (PLMs) have emerged as a promising area of study. PLMs are typically pre-trained on extensive sequence databases like UniRef^42^ before being applied to downstream tasks. This pre-training process usually involves masked language modeling, where parts of the input sequence are masked and the model is trained to predict the original amino acids in these masked positions. While this training process is relatively simple, conducting it on a large scale allows the model to understand relationships between amino acids, evolutionary information, and structural details present in protein sequences. This capability enables the model to efficiently extract useful information from the sequences^43^. For instance, PLMs have successfully uncovered implicit relationships among protein sequences, structures^38^, functional motifs, post-translational modifications^44^, and cellular localization^45^, thereby reducing the need for potentially laborious experimental investigations. Given the vast amount of labeled protein data from MS-based proteomics, including protein half-lives, which are challenging to process manually, PLMs are expected to facilitate a more accurate understanding of the relationship between protein sequences and half-lives.

Herein, we propose the Protein Lifetime Neural Model (PLTNUM), an AI platform designed for a sequence-dependent prediction of protein half-lives, leveraging PLMs. PLTNUM requires only amino acid sequences as input to predict protein half-lives, achieving higher prediction accuracy by incorporating structural information from AlphaFold2^46,47^. Additionally, PLTNUM can be used to explore degron candidates by explaining the output in terms of the contribution of input values.

## Results

Fig. 1a presents an overview of PLTNUM’s workflow. PLTNUM was built on SaProt^39^, a PLM that incorporates 3D structural information from many structures predicted by AlphaFold2. SaProt leverages Foldseek^48^ to translate 3D structures into token sequences, providing a straightforward method to integrate structural information into the input. Although SaProt’s model architecture is equivalent to ESM-2^43^, which excels in various downstream tasks related to protein structure and function prediction, SaProt outperforms ESM-2 by learning the relationship between sequence and structural information during pre-training. Here, we compared the performance of PLTNUM with that of long short-term memory (LSTM)^49^ and ESM-2 650M models, both with and without pre-training. Because LSTM, a type of recurrent neural network (RNN)^50^, has been reported to perform well in proteomics tasks such as protein fragmentation prediction, intensity prediction, and charge state prediction^51–53^, it is suitable for comparison with the other models.

**Fig. 1:**
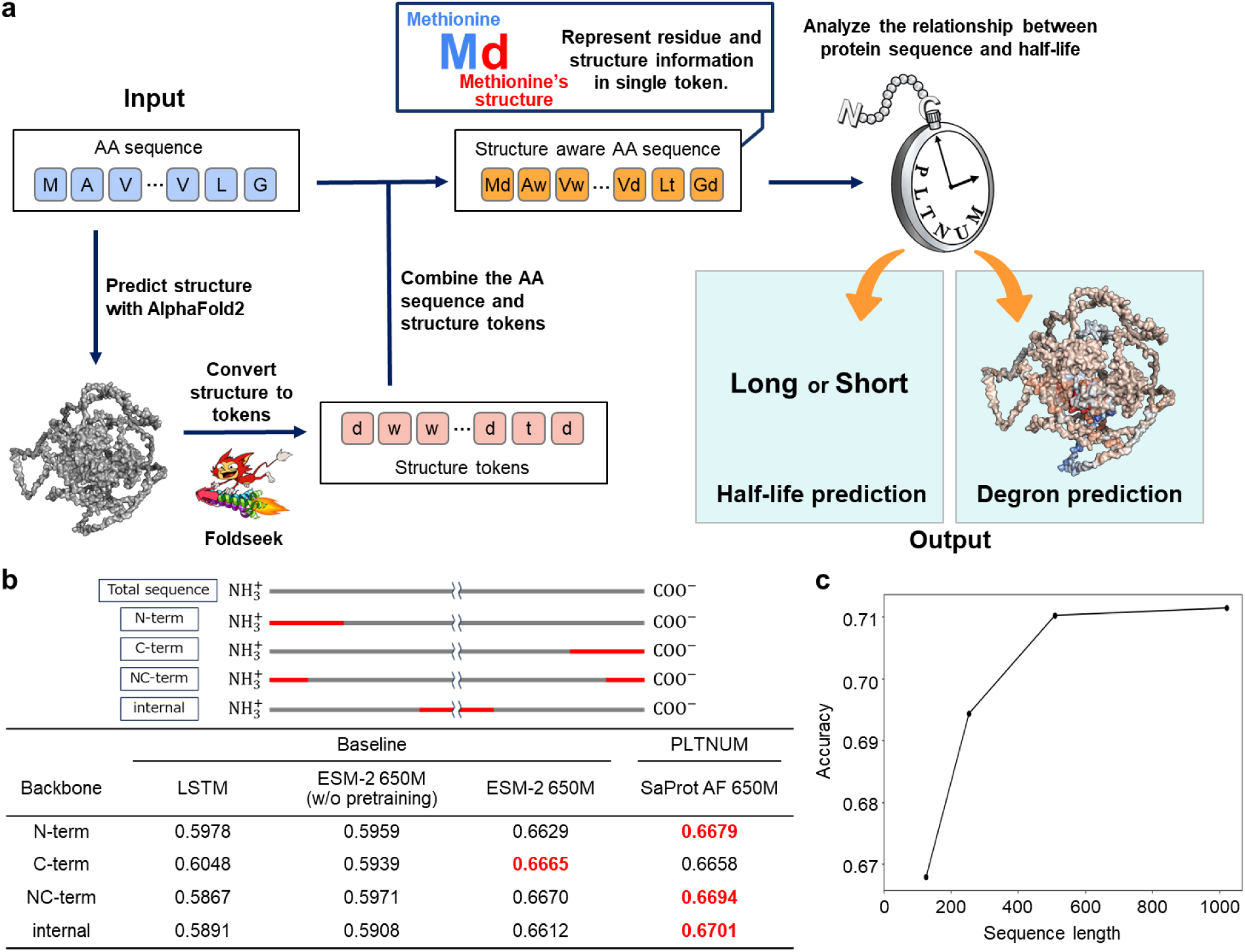
Basic Performance of PLTNUM in Protein Half-Life Prediction. **a**, Overview of PLTNUM’s workflow. **b,** Diagram of each segment used as input data and the accuracy of the tested models: LSTM, ESM-2 650M (without pre-training), ESM-2 650M (with pre-training), and PLTNUM. Each input sequence consists of 126 amino acid residues, and the accuracy of each model is summarized in the table. **c,** Plot of prediction accuracy versus input sequence length. The accuracy improved as the input sequence length increased, reaching a plateau at approximately 500 residues.

The dataset of protein half-lives from NIH3T3 mouse embryo fibroblast cell line^18^ was used to train PLTNUM and evaluate its predictive quality. When a peptide sequence detected by LC/MS/MS was present in multiple protein entries in the sequence database, the proteins were grouped, and the head protein was used as the representative. To incorporate protein 3D structural information as input data for PLTNUM, the dataset was narrowed down to 4,162 proteins registered in the AlphaFold Protein Structure Database^46,47^. To mitigate potential biases in the training dataset that could affect generalizability and model robustness, we investigated the distributions of protein half-lives and lengths within the dataset (Extended Data Fig. 1a, b). No discernible trends were found in these distributions. We also compared the biological functions of proteins in the training dataset with those in Swiss-Prot^54^ using the DAVID gene functional classification tool^55^ (Extended Data Fig. 1c, d). The high similarity in the top 10 gene ontology (GO) terms related to cellular components and biological processes between the two datasets suggested that the training data are not biologically biased.

PLTNUM predicted protein half-life with higher accuracy than the other models (Fig. 1b). The LSTM model showed low accuracy in predicting protein half-lives. Despite its more sophisticated architecture and 300 times more parameters, ESM-2 650M without pre-training exhibited similarly low accuracy to LSTM. However, ESM-2 650M with pre-training showed improved accuracy compared to its non-pre-trained counterpart, while SaProt AF 650M achieved even higher accuracy than the pre-trained ESM-2 650M. These results underscore the importance of pre-training on large datasets and incorporating structural information for the high prediction accuracy of protein half-lives by PLTNUM. The accuracy differences among input data from different protein positions were small; thus, the N-terminus was selected for subsequent experiments. Additionally, we investigated the dependence of accuracy on the N-terminus length of the input data (Fig. 1c). As the N-terminus length increased, the accuracy improved, reaching a plateau at approximately 500 residues and achieving a high prediction accuracy of 0.71. Considering both prediction accuracy and computational cost, the sequence length was fixed at 510 residues for subsequent experiments.

To understand how PLTNUM interprets protein half-lives, we visualized the <cls> token embedding representation output by the backbone language model in 2D space using t-distributed stochastic neighbor embedding (t-SNE)^56^. The proteins in the two-dimensional t-SNE representation were clustered into 15 groups using density-based spatial clustering of applications with noise (DBSCAN)^57^ (Fig. 2). Despite using only the N-terminus sequence as input data, PLTNUM demonstrated an ability to learn the biological background of proteins (Extended Data Table 1). For instance, in cluster 13, the proteins were predicted to be short-lived, and GO enrichment analysis indicated that many of these proteins were related to the cell cycle. Cell-cycle-related proteins have short half-lives to prevent incorrect regulation and enable the rapid removal of old or damaged proteins during the cell cycle^17^. Interestingly, although proteins in clusters 8, 9, and 10 were all labeled as nuclear-related, PLTNUM predicted that proteins in clusters 8 and 9 would be short-lived, while those in cluster 10, which included histone-related proteins, would be long-lived. Nuclear-related proteins are generally short-lived because they are involved in dynamic processes requiring quick responses and timely regulation, such as transcription and cell cycle progression^1,58^. However, histone proteins are long-lived to maintain chromatin structure, ensure consistent gene regulation, and preserve genetic information over time^59,60^. These results suggest that PLTNUM implicitly learned not only the localization but also the function of each protein, potentially contributing to a deeper understanding of the functional role of protein half-lives.

**Fig. 2:**
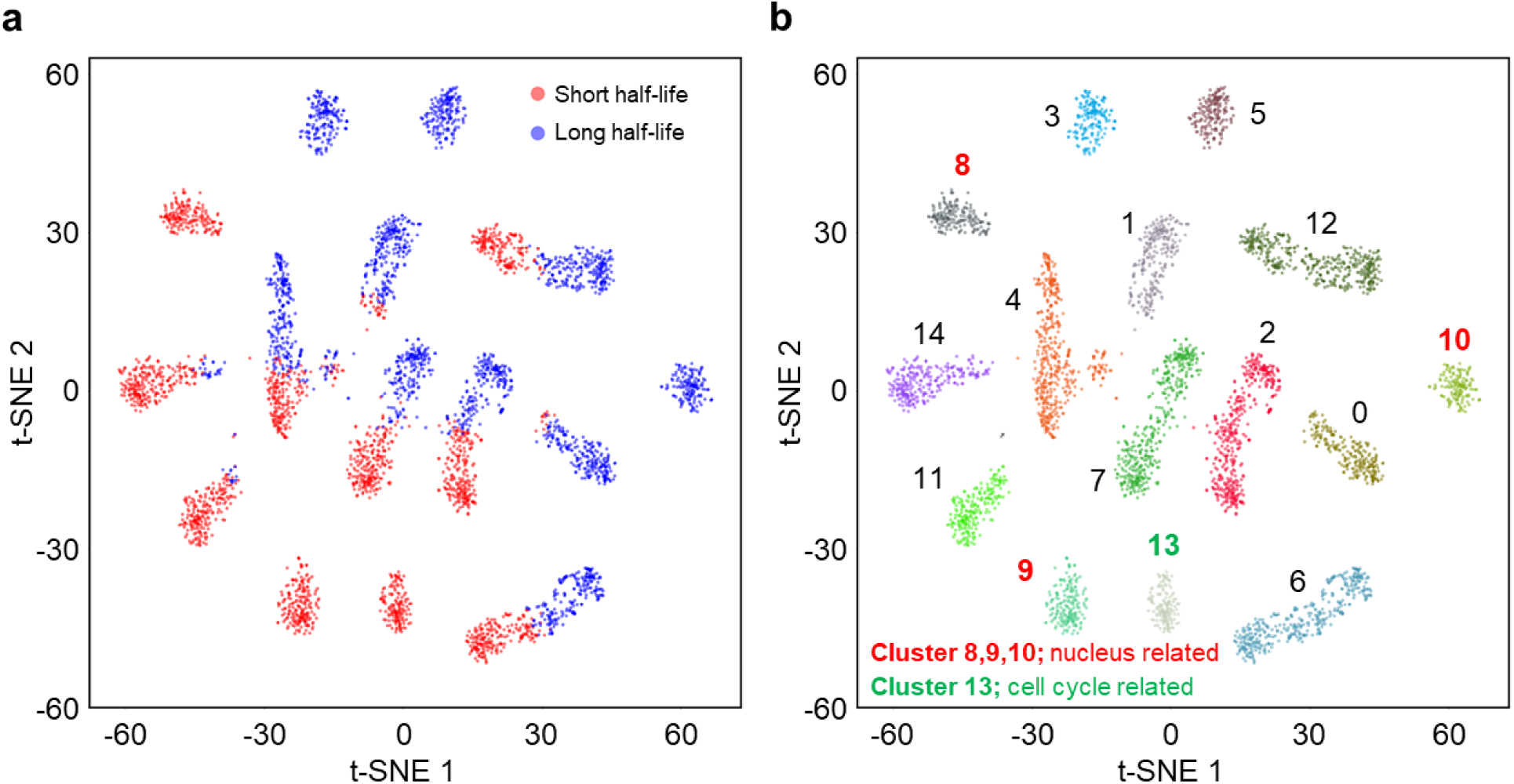
Visualization and clustering of protein half-life predictions by PLTNUM. **a,** Two-dimensional t-SNE representation of <cls> token embeddings output by the backbone language model. Proteins are colored based on their predicted half-lives: red for short lives and blue for long lives. **b,** Clustering of the t-SNE plot using DBSCAN, resulting in 15 clusters, each colored differently. GO enrichment analysis annotated each cluster as follows: clusters 0-7 are mitochondria-related, clusters 8-10 are nucleus-related, clusters 11 and 12 are PTM-related, cluster 13 is cell cycle-related, and cluster 14 is nucleotide binding-related.

One of the goals of developing PLTNUM is to understand the practical relationship between amino acid sequences and protein half-lives and to use it as a tool for identifying degron candidates. To achieve this, we utilized Shapley Additive Explanations (SHAP)^61^ values. SHAP is a well-known contribution calculation algorithm based on game theory, used to quantitatively evaluate the contribution of each input factor to the predicted output. In the context of PLTNUM, SHAP values quantify the contribution of each amino acid, including its type and position, to the difference between the model’s predicted value and a reference value of each protein, ensuring that the total contribution of all amino acids equals this difference. In brief, factors with positive SHAP values contribute to longer protein half-lives, while those with negative SHAP values contribute to shorter half-lives.

As a result, cysteine (C) was found to significantly contribute to the shortening of protein half-lives (Fig. 3a). The N-degron cysteine is known to trigger O2-dependent protein degradation through its oxidation and arginylation, leading to proteolysis by either the ubiquitin-proteasome system or autophagy^24^. This finding supports PLTNUM’s prediction that cysteine contributes to shorter protein half-lives. Furthermore, the average SHAP value of amino acids at each position from the N-terminus tends to be greater the closer they are to the end (Fig. 3b), supporting the N-end rule, which states that degrons are often located near the N-terminus^22^.

**Fig. 3:**
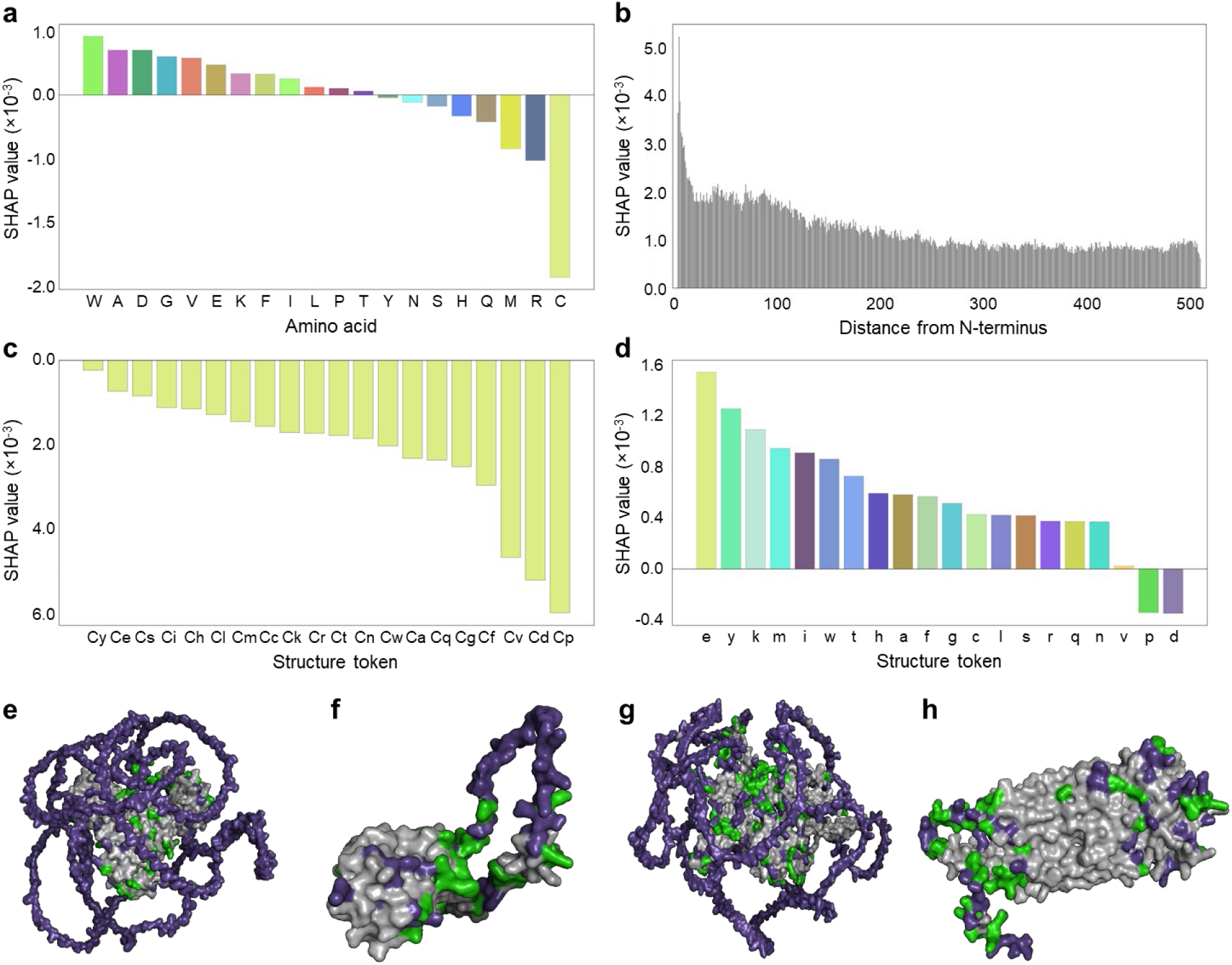
Contributions of amino acids, positions, and structural information to protein half-lives. **a,b,c,d,** SHAP values indicating the contribution of (a) amino acids, (b) position, and (c), (d) structure tokens. **e,f,g,h,** Visualization of structure tokens contributing to short protein half-life, as revealed by SHAP values. Color-coding indicates the concentration of structures “p” (green) and “d” (purple) in intrinsically disordered regions. (e) Q8R4X32, (f) Q9CPR1, (g) Q69ZW3, (h) Q6DID7. The three-dimensional structures of each protein were predicted by AlphaFold.

A standout feature of PLTNUM is its ability to evaluate the contribution of structural information of amino acids to protein half-lives, as PLTNUM incorporates structure tokens into the input sequence data using Foldseek. The contribution of cysteine, for example, varies significantly depending on the assigned structure token (Fig. 3b), suggesting that structural information plays a crucial role in degron functionality. Encouraged by this finding, we calculated the contribution of structure tokens for each amino acid (Extended Data Fig. 2). Comparing the average values of each structure token, we found that “p” and “d” tokens significantly contribute to shortening protein half-lives (Fig. 3d). Color-coding in the three-dimensional structures predicted by AlphaFold^47^indicates that these structure tokens are concentrated in the intrinsically disordered regions on the exterior of proteins (Fig. 3e, f, g, h). This result aligns with previous reports that degrons often reside in intrinsically disordered regions^6,17,62,63^. In this way, PLTNUM can quantitatively evaluate the contributions of amino acid types, positions, and structural information to protein half-lives, providing numerous insights for predicting unknown degrons.

One of the significant advantages of PLTNUM is that it uses only protein sequence information and half-life data for training, relying on a comprehensive protein half-life dataset. This approach enables unbiased degron predictions, unlike previous models that were trained on reported degron sequences and thus might produce biased outcomes^6,62,64^. For example, the half-life of individual proteins is influenced by a complex interplay of factors that either shorten or lengthen it. However, traditionally known degron sequences have been identified primarily from short-lived proteins, with long-lived proteins overlooked in many studies. In our study, we calculated the SHAP values of sequences consisting of 1 to 5 amino acids, identifying the top 10 combinations with the most negative SHAP values as potential degron sequences (Fig. 4b). To minimize noise in the dataset and ensure the robustness of our method, we considered only sequences that appeared more than five times in the protein sequences of the training dataset. To evaluate the prediction accuracy of these potential degron sequences, we compared the average half-life of proteins containing these sequences (Group A) to that of proteins with the same composition but different sequence orders, including structure tokens (Group B) (Fig. 4a). The results showed that Group A had shorter protein half-lives for 8 out of the top 10 candidate degron sequences compared to Group B, indicating that the degron prediction by PLTNUM is highly reliable. Interestingly, the top 8 sequences consistently contained a cysteine residue and tended to shorten the protein half-lives. In line with many reports, this result implies the importance of both the presence and position of cysteine residues in degron sequences.

**Fig. 4:**
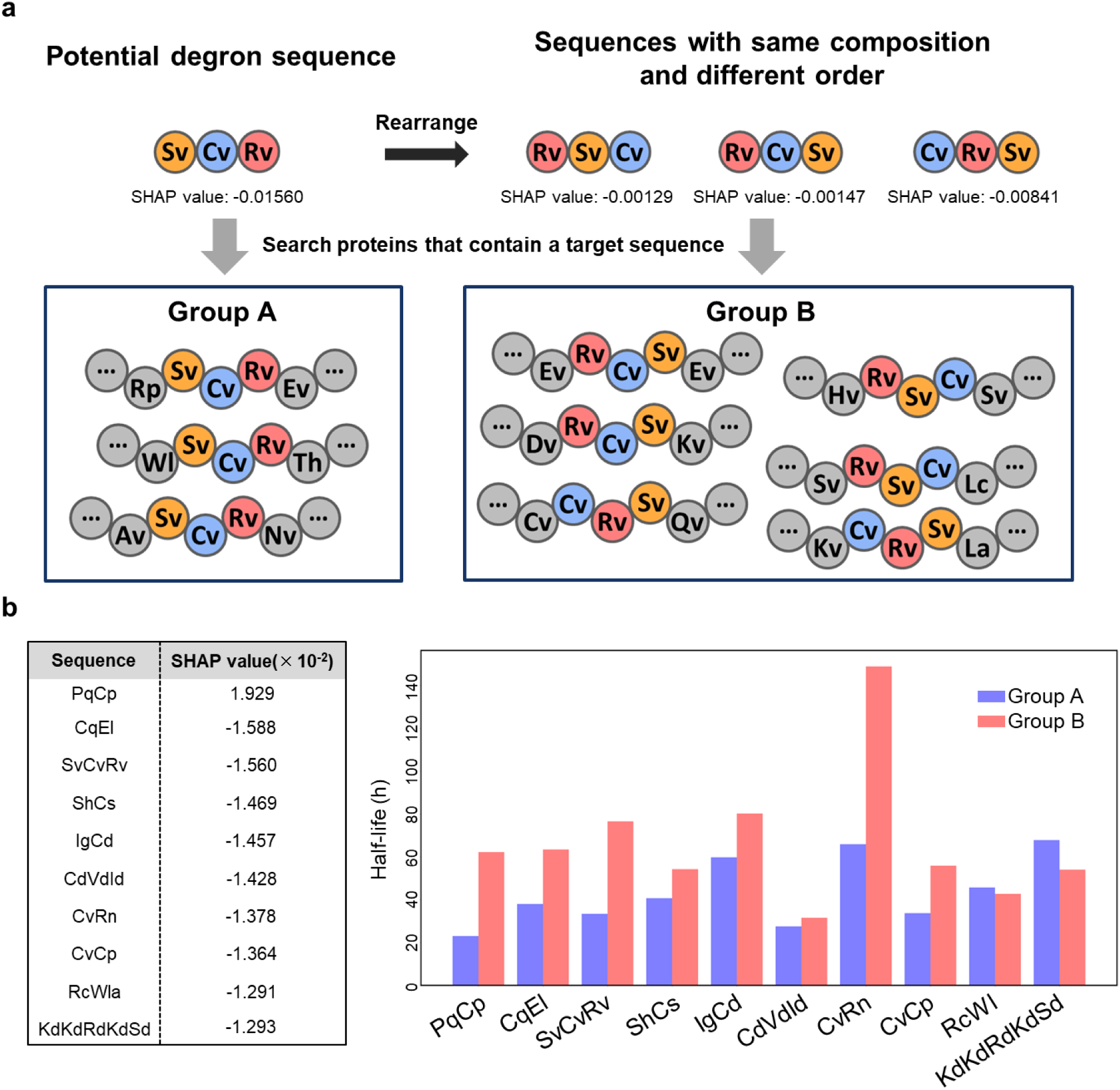
Identification of potential degron sequences using PLTNUM. **a,** Workflow for evaluating PLTNUM’s prediction accuracy for potential degron sequences. Potential degron sequences consisted of 1 to 5 amino acids with top 10 negative SHAP values. Group A included proteins with these potential degron sequences, while Group B included proteins with sequences having the same amino acid composition as the potential degron but in a different order. The prediction accuracy was evaluated by comparing the average half-lives of these groups. **b,** Average half-lives of Group A and Group B. The left column of the left table shows the potential degron sequences, and the right column shows the SHAP values.

To further confirm the high half-life prediction accuracy of PLTNUM, we compared its performance with that of existing protein stability prediction models (Fig. 5a, b). DEGRONOPEDIA^64^ predicts the protein stability index (PSI), which is calculated based on global protein stability data, explicitly targeting the stability of terminal peptides. The coefficients of determination are 0.812 for N-terminus with methionine, 0.796 for N-terminus without methionine, and 0.815 for C-terminus. For comparison, we used the model that predicts PSI for N-terminus with methionine. We also used TemStaPro^65^ and DeepSTABp^66^ for comparison; these models were developed to predict protein thermal stability, and can accurately predict melting points based on sequence information (AUC = 0.993, R^2^ = 0.80, respectively). We applied these models to the dataset of proteins in NIH3T3 cells, which was used to train PLTNUM, and created receiver operating characteristic (ROC) curves (Fig. 5a). The ground truth labels were determined based on whether the protein half-life was greater or less than the median of the dataset. PLTNUM showed superior performance in predicting protein half-life compared to the other models (Fig. 5a). Furthermore, PLTNUM demonstrated high predictive performance not only on the NIH3T3 dataset used for training but also on a dataset of HeLa cells measured under entirely different conditions^19^ (Fig. 5b), indicating its high generalizability. In summary, PLTNUM advances protein half-life prediction using only amino acid sequences while incorporating structural information, showcasing robustness and versatility across different datasets and experimental conditions.

**Fig. 5:**
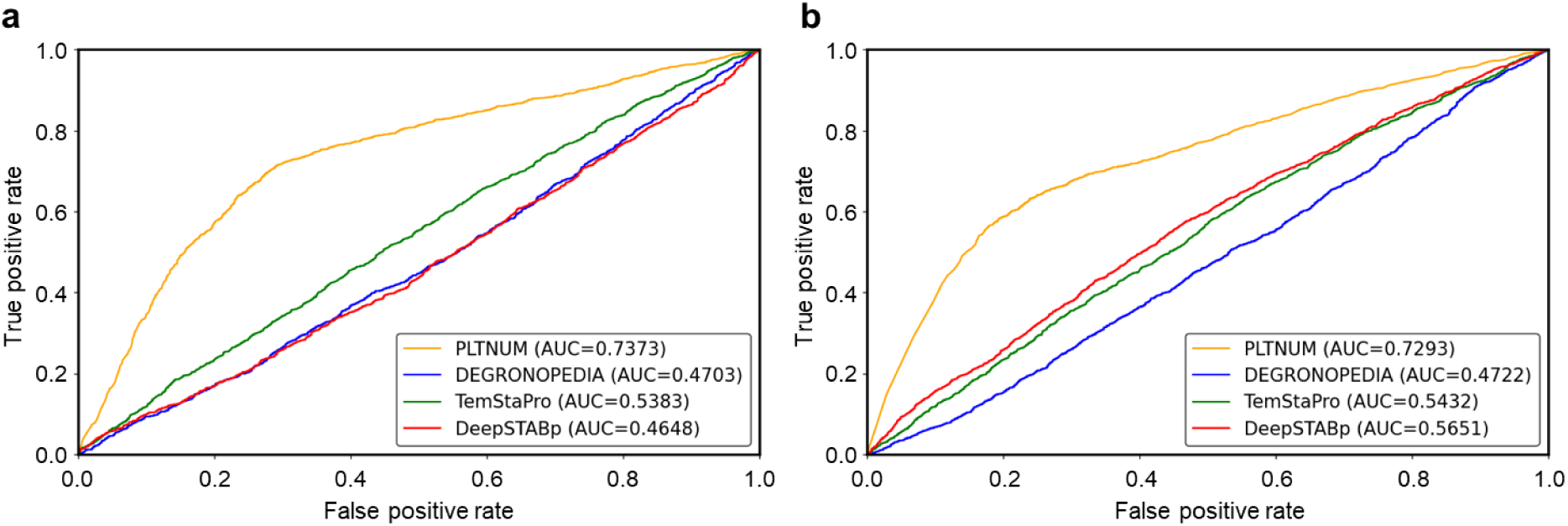
Performance comparison of PLTNUM and existing protein stability prediction models for protein half-life prediction. **a,** ROC curves for PLTNUM, DEGRONOPEDIA, TemStaPro, and DeepSTABp applied to a protein half-life dataset from NIH3T3 cells. The area under the curve (AUC) values are shown in the legend, and indicate that PLTNUM (AUC = 0.7373) outperforms the other models (DEGRONOPEDIA AUC = 0.4703, TemStaPro AUC = 0.5383, DeepSTABp AUC = 0.4648). **b,** ROC curves for the same models applied to a protein half-life dataset from HeLa cells. The AUC values are shown in the legend, indicating PLTNUM (AUC = 0.7293) outperforming DEGRONOPEDIA (AUC = 0.4722), TemStaPro (AUC = 0.5432), and DeepSTABp (AUC = 0.5651).

## Discussion

PLTNUM, the first protein half-life prediction model that leverages protein language models and comprehensive half-life datasets, significantly outperforms existing protein stability prediction models, such as DEGRONOPEDIA, TemStaPro, and DeepSTABp. Protein half-lives are determined by a complex interplay of sequence-dependent factors, such as amino acid sequences and structures, and biological context-dependent factors, such as species, organs, cell types, and organelles. While PLTNUM primarily considers sequence-dependent factors, it has also been shown to implicitly learn biological context-dependent factors, such as protein localization and function. We confirmed its generalizability across various measurement conditions and cell types.

PLTNUM can also be directly applied to predict the half-lives of mutant sequences. Given the association between unusually long protein half-lives and diseases such as cancer, PLTNUM appears to have the potential to serve as a screening tool to identify disease-causing mutations. Our study also evaluated the impact of protein structure on half-life prediction. We found that incorporation of SaProt, which converts 3D structural information of proteins into a string format, improved the predictive accuracy of PLTNUM. This finding emphasizes the importance of considering protein structural information in protein half-life prediction and confirms that structural factors influence protein stability.

A notable contribution of this study is the quantitative investigation of the contribution of each amino acid, its order, and structural information to protein half-lives, facilitated by the integration of PLTNUM and SHAP. Unlike methods that rely on already known degron sequences, our approach predicts degron sequence candidates from scratch, thus avoiding inherent biases. PLTNUM revealed that cysteine and intrinsically disordered regions contribute to shortening the half-life of proteins, identifying potential degron sequences that may contribute to this trait. Our validation study confirmed that proteins containing these candidate degron sequences generally have shorter half-lives, supporting the effectiveness of our method for comprehensively identifying degron sequences.

In conclusion, PLTNUM is the first model to predict protein half-lives from amino-acid sequences. While PLTNUM can predict degrons in an unbiased manner, its prediction accuracy remains at only approximately 71%, emphasizing the need to consider the biological context in addition to the sequence information for further improvement. From another perspective, PLTNUM is also expected to be utilized as a filter for exploring proteins whose half-lives are influenced by biological context.

## Methods

### Model Construction

PLTNUM was constructed based on SaProt AF 650M^39^. To benchmark the performance, we constructed other protein half-life prediction models, including the LSTM model^49^ and the ESM-2 model, both with and without pretraining^43^. All models and training codes were implemented in Python (version 3.11), using libraries such as Pytorch (version 2.3.1)^67^, Transformers (version 4.38.2)^68^, and Scikit-Learn (version 1.2.2)^69^. The architecture of the LSTM model included two layers of bidirectional LSTM with 257 nodes each, followed by a dropout layer^70^ and a fully connected layer. PLTNUM and ESM-2 models utilized PLM encoders from the Transformers library, incorporating dropout layers and a fully connected layer. The output of all models was a single-dimensional value, scaled between 0 and 1 using the sigmoid function.

We applied a high dropout rate of 0.8 for all models to mitigate overfitting and improve generalizability. Additionally, multi-sample dropout was employed in the PLM-based models to further reduce overfitting. For the LSTM model, input sequences were one-hot encoded, while the PLM-based models used tokenized sequences created by their respective tokenizers. We standardized the input sequence length to ensure uniformity across the dataset: sequences shorter than the specified length were padded, while longer sequences were truncated. Tokenized sequences for PLM-based models included a <cls> token at the beginning and an <eos> token at the end. When utilizing NC-term as input, N-terminal and C-terminal sequences were concatenated with a space to differentiate the information.

The model outputs were continuous values, which were binarized using a threshold of 0.5. Outputs below this threshold were classified as “0” (short-lived), while those above were classified as “1” (long-lived). This threshold was chosen as a standard for binary classification to ensure clear differentiation between short-lived and long-lived proteins.

### Model Training and Evaluation

We used a 10-fold cross-validation approach for model training and evaluation. The LSTM model was trained with a learning rate of 0.0002, while the PLM-based models were trained with a learning rate of 0.00002. All models were trained for 10 epochs in a supervised manner, using binary cross-entropy loss as the objective function. To prevent overfitting in the PLM-based models, we applied a data augmentation technique where 5% of the tokens were randomly replaced with <mask> tokens with a probability of 20%. This technique was designed to encourage the models to better understand sequence contexts, thereby enhancing their generalizability.

During training, model performance was monitored using the F1 score on the validation data. The checkpoint corresponding to the epoch that achieved the highest F1 score was saved for each model. For the final evaluation, inference was performed on the validation data for each fold. The accuracy of the models was calculated after combining the validation data across all folds, providing a comprehensive assessment of each model’s performance across the entire dataset.

### Visualization of PLTNUM’s Output

We visualized PLTNUM’s internal representations to gain insight into how it interprets protein half-life. After training PLTNUM using 10-fold cross-validation, we input the validation data for each fold into the model to extract the <cls> token embeddings from the PLTNUM’s backbone PLM, SaProt. These embeddings, which are 1280-dimensional vectors, encapsulate comprehensive information about the protein sequences and half-lives.

We reduced these high-dimensional vectors to two dimensions using t-SNE and subsequently performed clustering with the DBSCAN algorithm. DBSCAN was chosen for its robustness in detecting clusters of arbitrary shapes and handling outliers without requiring a predefined number of clusters. For our analysis, we set the hyperparameter epsilon to 0.5 and the minimum number of points per cluster to 5. This resulted in the formation of 15 clusters, with three proteins classified as outliers.

To interpret the biological significance of these clusters, we conducted enrichment analysis using the DAVID gene functional classification tool. The results of this analysis are presented in Extended Data Table 1.

### Quantification of Amino Acid Contributions to Protein Half-lives

SHAP^61^ values were used to quantify the contribution of each amino acid and structural information to PLTNUM’s protein half-life predictions. For each validation dataset in the 10-fold cross-validation, we used the trained PLTNUM model to compute SHAP values. These were calculated using the partition explainer from the SHAP Python library (version 0.45.1), with the “max_evals” parameter set to 5000. SHAP values were determined per token, with each token representing a combination of amino acid and its structural information. The same SHAP value was assigned to both the amino acid and structure token to evaluate the contribution of amino acid or structural information separately.

The average SHAP value for each amino acid was calculated by averaging the SHAP values across all occurrences of that amino acid in the dataset. The average SHAP value for each structure token was obtained by averaging the SHAP values of all instances of that structure token. Additionally, to evaluate the impact of residue position relative to the N-terminus on protein half-life prediction, SHAP values were calculated for all proteins. The average absolute SHAP values for each position from the N-terminus were then calculated to determine the contribution of the positioning to model predictions.

### Comparison of PLTNUM’s Prediction Accuracy with Reported Model

We compared the protein half-life prediction accuracy of PLTNUM with reported protein stability prediction models, including DEGRONOPEDIA^64^, TemStaPro^65^, and DeepSTABp^66^. DEGRONOPEDIA employs CatBoost regression models^71^ to predict the protein stability index (PSI), derived from global protein stability experiments. For this comparison, we used the DEGRONOPEDIA model that predicts the PSI of N-terminal 23-mers with an initiator methionine. TemStaPro utilizes protein embeddings from the ProtT5-XL model^72^ to predict protein stability at specific temperature thresholds. We used the TemStaPro-40 model, which is trained with a 40 °C threshold and demonstrated the highest AUC in protein half-life prediction. DeepSTABp predicts the melting temperature of proteins using embeddings from ProtT5-XL-UniRef50. Predictions were made for proteins in a cell with a growth temperature of 37 °C.

We applied PLTNUM and the three comparative models to protein datasets from NIH3T3^18^ and HeLa cells^19^. Proteins were classified as “long-lived” or “short-lived” based on the median half-life within each dataset. Predictive performance was evaluated using receiver operating characteristic (ROC) curves. For the NIH3T3 dataset, predictions were made on the validation data in a 10-fold cross-validation setting. For the HeLa dataset, predictions were made using one of the ten PLTNUM models generated during cross-validation.

## Acknowledgements

We would also like to thank our lab members in Kyoto University for fruitful discussions. This work was supported by JSPS Grants-in-Aid for Scientific Research (23K13774, 21K14652, 20K20567 to E.K., 23H04924, 23K18185, 21H02459 to Y.I.), JST Strategic Basic Research Program CREST (JPMJCR1862), AMED-CREST program (JP18gm1010010) and NEDO Intensive Support Program for Young Promising Researchers (seeds-4891) to E.K.

## Author information

Authors and Affiliations

**Graduate School of Pharmaceutical Sciences, Kyoto University, Kyoto, Japan**

Tatsuya Sagawa, Eisuke Kanao, Kosuke Ogata, Koshi Imami & Yasushi Ishihama

**Laboratory of Proteomics for Drug Discovery, National Institute of Biomedical Innovation, Health and Nutrition, Ibaraki, Osaka, Japan**

Tatsuya Sagawa, Eisuke Kanao & Yasushi Ishihama

**Proteome Homeostasis Research Unit, RIKEN Center for Integrative Medical Sciences, Yokohama, Japan**

Koshi Imami

## Contributions

Y.I., K.I. and K.O. developed the initial concept. Y.I. provided resources for the study. T.S. designed and executed all computational experiments and data analyses. T.S. drafted the manuscript and designed the figures. E.K. and Y.I. supervised the work. K.I. and K.O. provided critical insights in analyses. Writing—review and editing was carried out by T.S., E.K. and Y.I. Approval of the final version of the paper was carried out by all authors.

## Ethics declarations

All authors declare no competing interests.

## Additional information

Extended data Fig. 1, Fig. 2 and Table 1 are available for this paper at XXX.

## Data availability

The original protein half-life datasets of NIH3T3 cell line and HeLa cell line are available in the Supplemental Table S3 of the paper by Schwanhäusser, B. *et al.*^18^ and in the Supplementary Material of the paper by Zacha et al.^19^, respectively.

## Code availability

The source code of PLTNUM is available on GitHub at https://github.com/sagawatatsuya/PLTNUM.

**Correspondence and requests for materials** should be addressed to Yasushi Ishihama.

## Table of contents

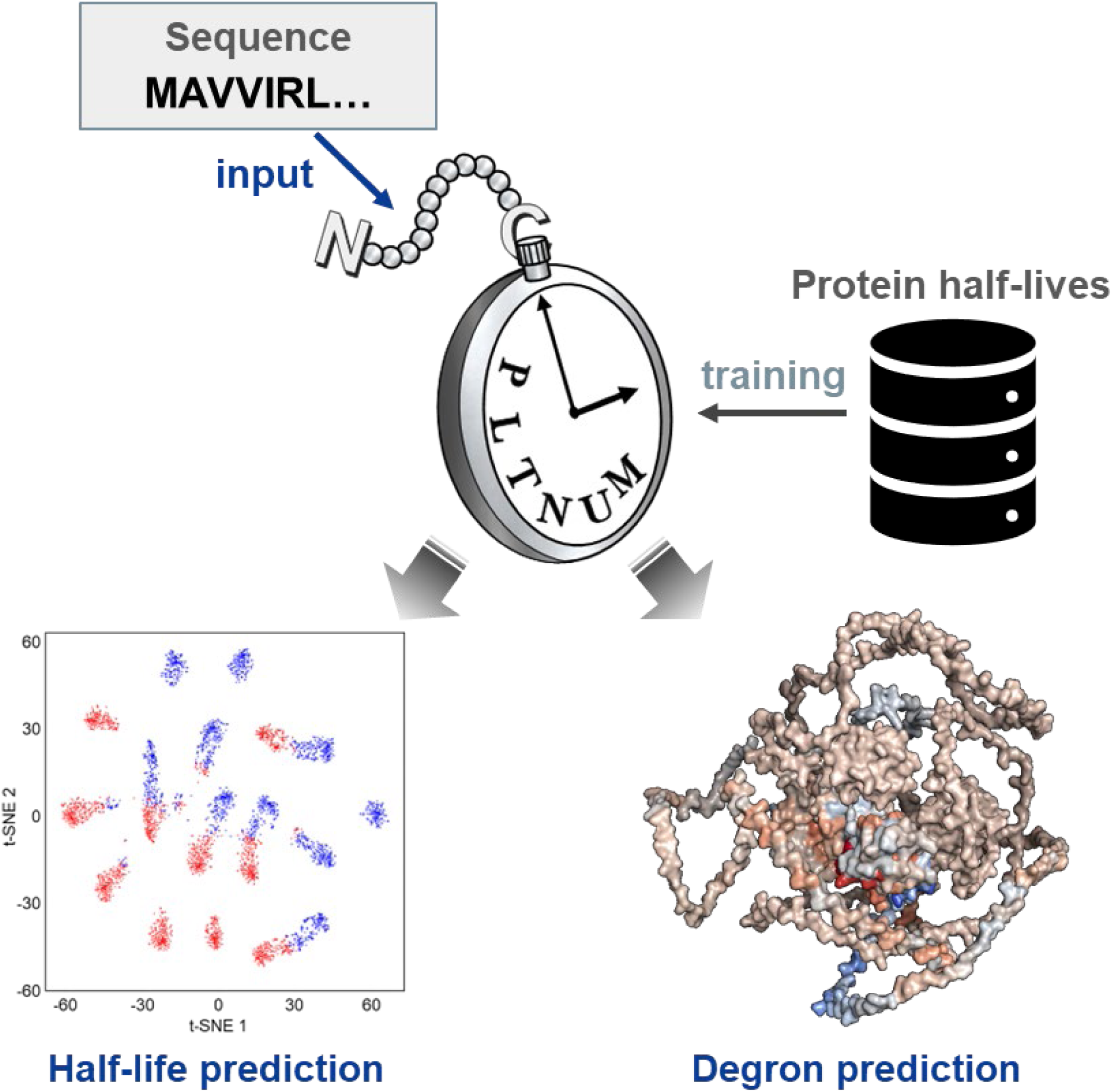

## Extended Data

**Extended Data Fig. 1.**
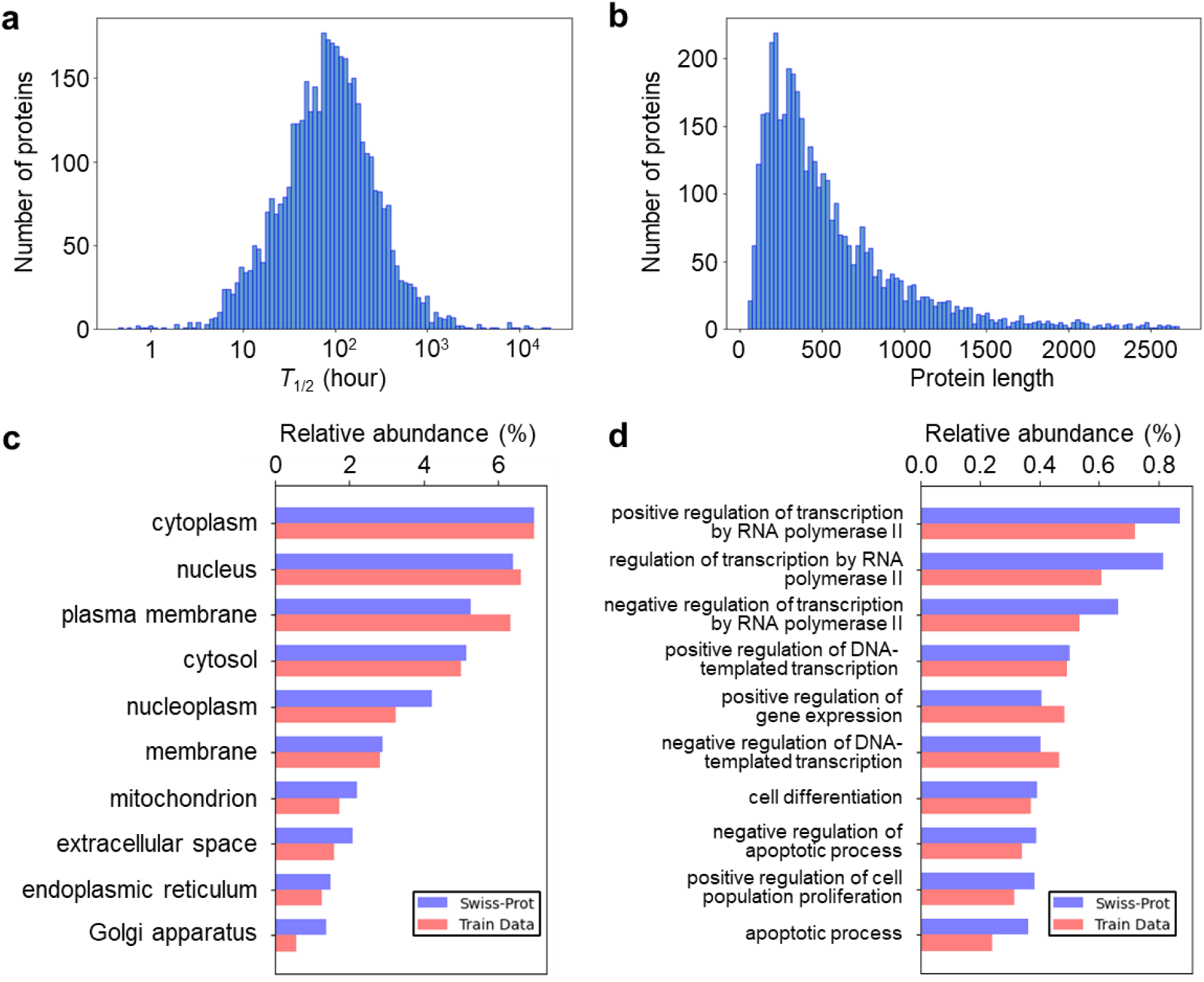
Features in the training dataset. 4,162 proteins registered in both the report by Schwanhäusser, et al.^18^ and the AlphaFold Protein Structure Database^46,47^ were used as the training dataset. a,b, Histograms of protein half-lives (a) and protein lengths (b) in the training dataset. c,d, Comparison of cellular components (c) and biological processes (b) between Swiss-Prot (blue bar) and training data (red bar). Each Gene Ontology term was obtained from UniProt database using David 6.8 Bioinformatics Resources.^55^

**Extended Data Fig. 2.**
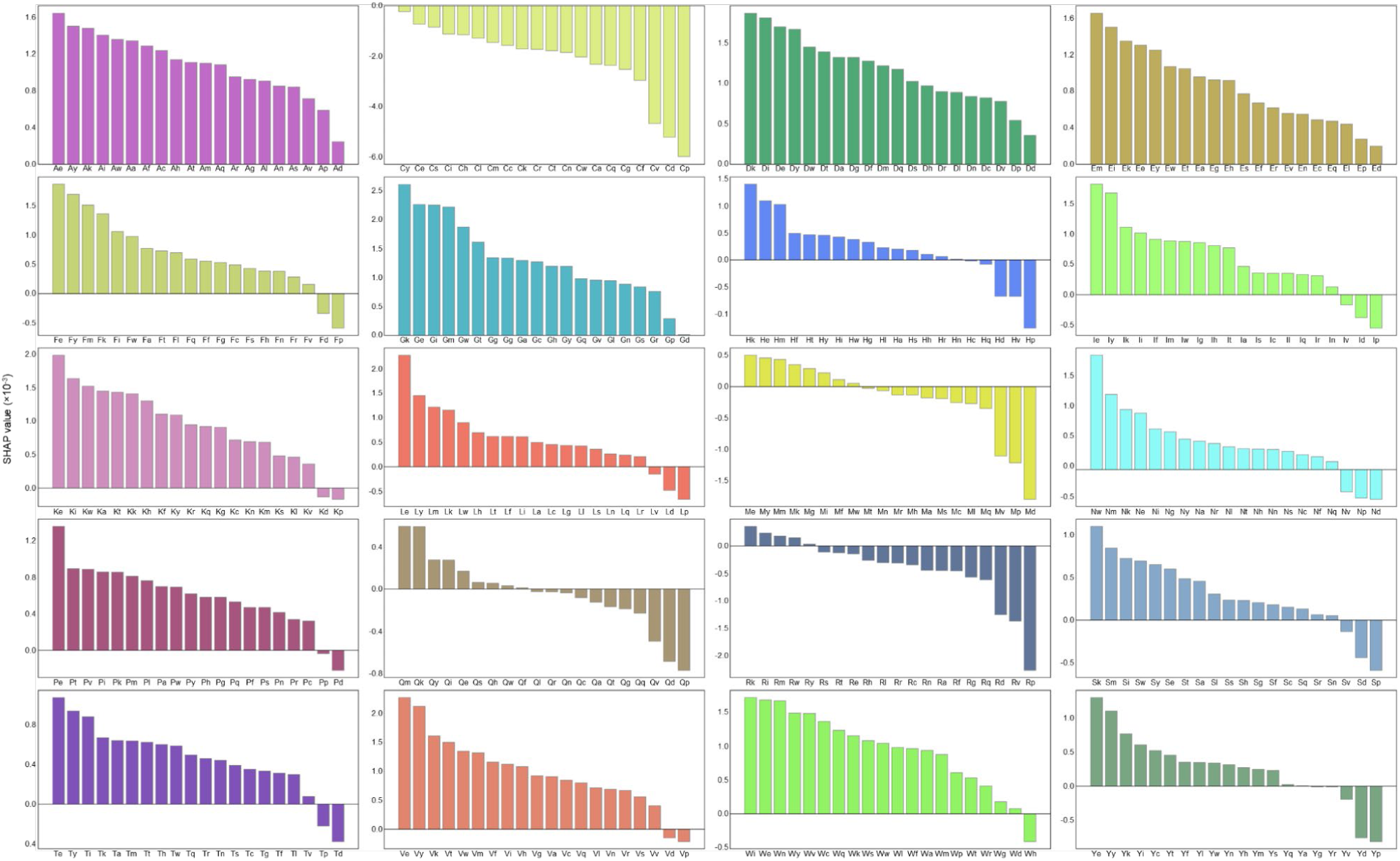
Contribution analysis of each amino acid to protein half-lives. The same amino acid showed various contributions to protein half-lives based on the structure.

---

**Extended Data Table. 1.**
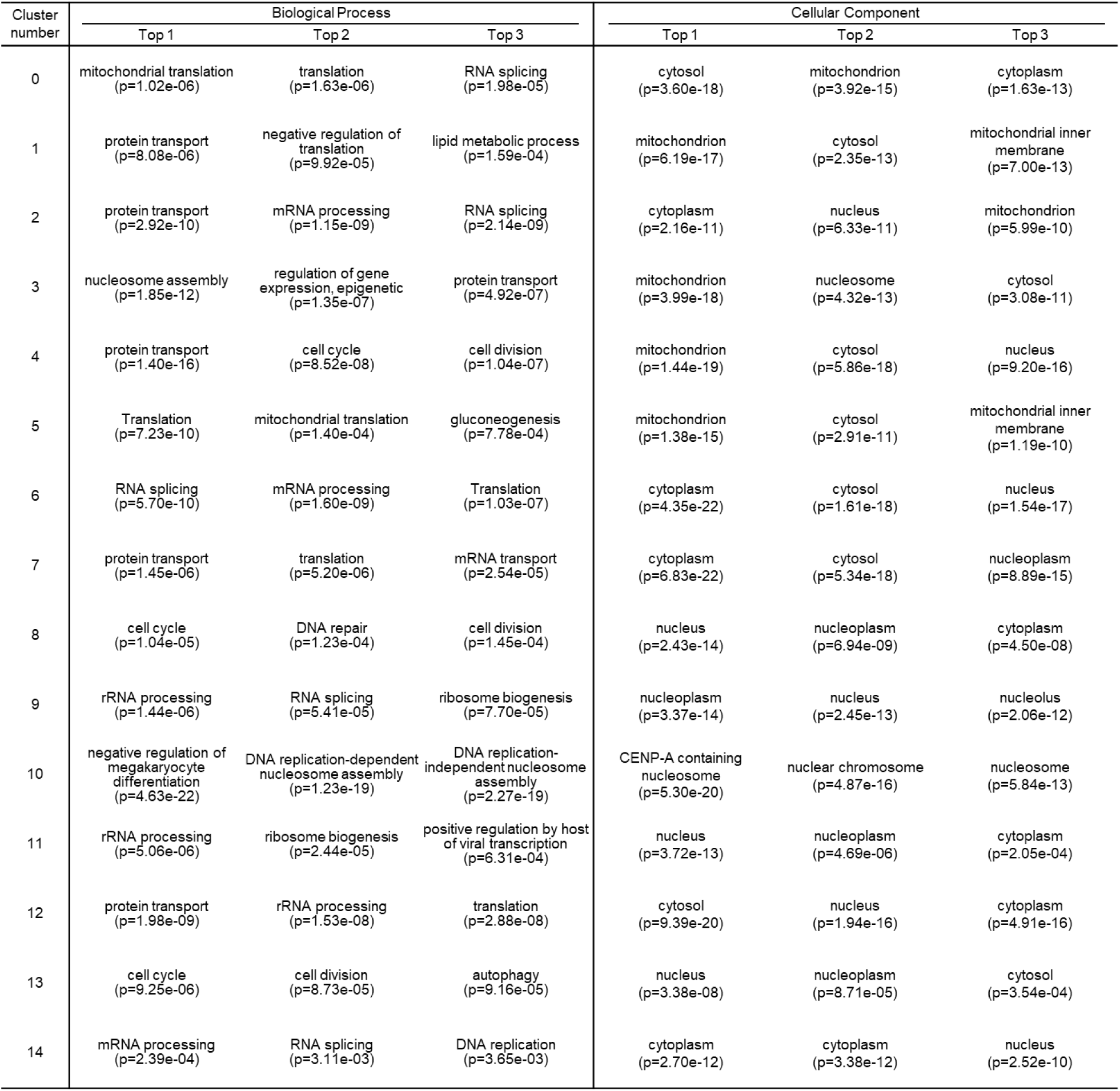
Clusters generated by PLTNUM. The result of GO enrichment analysis using the DAVID gene functional classification tool of 15 clusters generated by PLTNUM. In some clusters, significant enrichment of biological features was observed.

